# Vestibular prepulse inhibition of the human blink reflex

**DOI:** 10.1101/2024.01.09.574842

**Authors:** Matteo Ciocca, Sarah Hosli, Zaeem Hadi, Mohammad Mahmud, Yen Tai, Barry M Seemungal

**Author notes:** **Corresponding author:** Matteo Ciocca. **Present address:** Department of Brain Sciences, Imperial College London, Charing Cross campus, Fulham Palace Rd, London W6 8RF. **Authorship statement** All co-authors have reviewed and approved the contents of the manuscript, and the submission is not under review at any other publication.

## Abstract

**Objective:** Auditory and somatosensory prepulses are commonly used to assess prepulse inhibition (PPI). The effect of a vestibular prepulse upon blink reflex (BR) excitability has not been hitherto assessed.

**Methods:** Twenty-two healthy subjects and two patients with bilateral peripheral vestibular failure took part in the study. Whole body yaw rotation in the dark provided a vestibular inertial prepulse. BR was electrically evoked after the end of the rotation. The area-under-the-curve (area) of the BR responses (R1, R2, and R2c) was recorded and analysed.

**Results:** A vestibular prepulse inhibited the R2 (p < 0.001) and R2c area (p < 0.05). Increasing the angular acceleration did not increase the R2/R2c inhibition (p>0.05). Voluntary suppression of the vestibular-ocular reflex did not affect the magnitude of inhibition (p>0.05). Patients with peripheral vestibular failure did not show any inhibition.

**Conclusions:** Our data support a vestibular-gating mechanism in humans.

**Significance:** The main brainstem nucleus mediating PPI – the pedunculopontine nucleus (PPN) – is heavily vestibular responsive, which is consistent with our findings of a vestibular-mediated PPI. Our technique may be used to interrogate the fidelity of brain circuits mediating vestibular-related PPN functions. Given the PPN’s importance in human postural control, our technique may also provide a neurophysiological biomarker of balance.

**Highlights:**

- This is the first report of a vestibular prepulse inhibition of the blink reflex.
- A vestibular prepulse inhibits the R2/R2c area in healthy subjects but not in patients with bilateral peripheral vestibular failure.
- Vestibular PPI is a potential neurophysiological marker of vestibular-motor integration at the brainstem level.

## 1. Introduction

Sensorimotor gating is an essential central nervous system mechanism which allows the organism to focus on relevant stimuli and prevent overload or hyperresponsiveness to innocuous sensory inputs (Garcia-Rill et al., 2019, Kofler et al., 2023). Disruption of sensorimotor gating has been described in neurodegenerative disease, such as Parkinson’s disease (Millian-Morell et al., 2018), multiple system atrophy (Zoetmulder et al., 2014), and Huntington’s disease (Valls-Sole et al., 2004), and functional movement disorders (Hanzlikova et al., 2019).

Sensorimotor gating can be probed via prepulse inhibition (PPI), a neurophysiological technique in which a weak sensory stimulus results in inhibition of a subsequent test response (Gomez-Nieto et al., 2020). Theoretically, all sensory modalities may evoke an inhibition. Auditory and somatosensory prepulses are most commonly used in PPI studies, although a few have used visual and laser prepulse stimuli (Garcia-Rill et al., 2019). Whatever the sensory modality, the degree of inhibition depends upon both the intensity of the prepulse (higher intensities eliciting more inhibition) (Csomor et al., 2005), and timing of the prepulse, with optimal interstimulus intervals ranging from a minimum of 60 ms for acoustic to 800 ms for vibrotactile stimuli (Blumenthal, 2015). Gender and hormonal status also contribute to modulating PPI (Kumari et al., 2010, Naysmith et al., 2022).

PPI is mediated by an extensive network, with components including the basal ganglia, the cerebellum, limbic cortical inputs and the pedunculopontine nucleus (PPN), which appears to be a critical hub in mediating PPI since PPN lesions in animal studies abolishes PPI (Garcia-Rill et al., 2019). Interestingly, in animal models (including primates), the PPN receives significant innervation from the vestibular nuclei (Aravamuthan and Angelaki, 2012, Horowitz et al., 2005, Woolf and Butcher, 1989). Human neuroimaging and neurophysiological studies also support the notion that the PPN is part of the central vestibular network (Kirsch et al., 2016, Yousif et al., 2016).

Given the role of PPN in vestibular processing and its involvement in PPI, we hypothesized that vestibular stimuli are also filtered through such a gating mechanism via PPN and hence predicts the existence of a vestibular PPI, i.e. a weak activation of the vestibular system (i.e. a vestibular prepulse) would reduce the magnitude of a subsequent test response. A neurophysiological interrogation of vestibular-linked PPN activity has potential clinical utility given that imbalance and falls in Parkinson’s disease patients is linked to the loss of PPN cholinergic neurons (Karachi et al., 2010), and conversely PPN DBS modulates postural control in Parkinson’s Disease patients(Yousif et al., 2016).

To test our prediction of a vestibular PPI in healthy subjects, we combined blink reflex testing (via electrical stimulus to the supraorbital nerve) with a preceeding vestibular stimulus provided via whole-body, constant yaw-plane accelerations in the dark. If the vestibular pre-pulse inertial stimulus behaves congruently with standard PPI responses as seen with non-vestibular modalities (e.g. somatosensory), we predict a prepulse-associated suppression of the R2 response (figure 5 C-F), and a facilitation of the R1 response. To confirm the vestibular origin of our observations, we also assessed responses in patients with bilateral peripheral vestibular dysfunction. Vestibular failure patients, who lack a vestibular prepulse input to the brainstem circuitry responsible for PPI, would be predicted to show no vestibular PPI.

As for auditory and somatosensory PPI, we similarly hypothesised an intensity-dependent inhibition of the prepulse on the blink reflex R2 response (Blumenthal, 2015). Therefore, we predicted that the magnitude of suppression of the blink response will increase by increasing the intensity of the vestibular prepulse as defined by the inertial angular acceleration. To support our hypothesis, we tested four different prepulse angular accelerations.

## 2. Methods

### 2.1. Participants

We designed three separate experiments. Fifteen subjects (6 women; aged 21-40) participated in Experiment 1, while 19 subjects (7 women; aged 21-40) participated in Experiment 2. Nine subjects participated in both experiments. Two patients clinically diagnosed with a bilateral vestibular failure participated in Experiment 3. Experiments were approved by the local Research Ethics Committee and were performed in accordance with the Declaration of Helsinki.

### 2.2. Experimental set-up

We activated the vestibular system by using a passive, yaw-axis, whole-body rotation in dark (Seemungal et al., 2004). Participants sat on a computer-controlled, motorised and vibration-free rotating chair in total darkness and were instructed to keep looking ahead, keeping their head still against the headrest, relax their face, and minimise any spontaneous blinking response.

We used a symmetric triangular angular velocity waveform, with constant accelerations of 5, 10, 20, or 30 degrees/second^2^ for one second and a deceleration of identical magnitude and duration but in the opposite direction. Thus, the triangular angular velocity profile lasted a total of 2 seconds (Figure 1).

**Figure 1:**
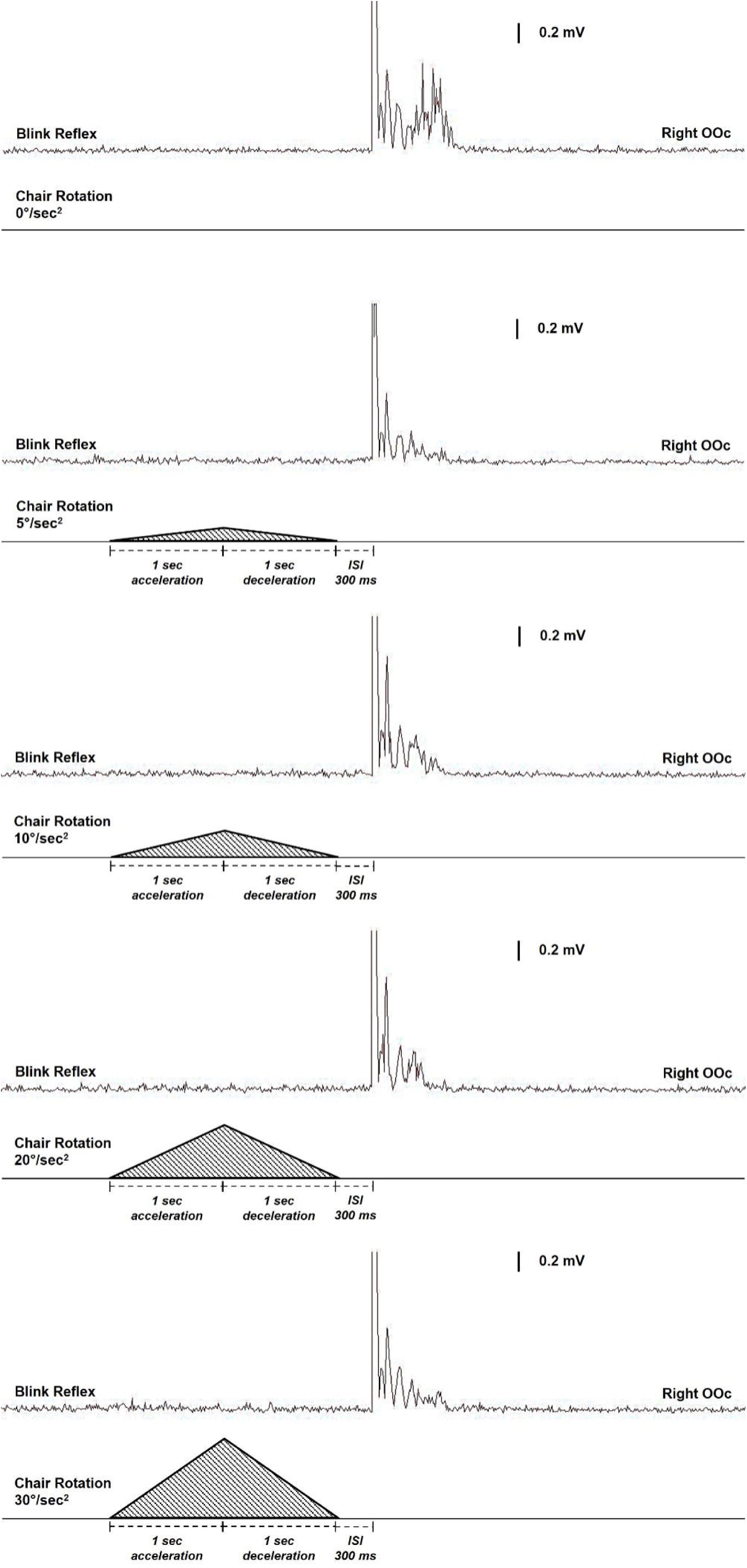
experimental set-up and prepulse modulation of the electrically induced R2 response of the blink reflex in one representative subject. The four different angular accelerations are reported, together with the rectified averaged EMG trace from the right orbicularis oculi for each condition. An inhibition of the R2 response and a facilitation of the R1 response are present in this subject.

Three hundred milliseconds after the end of the rotation, we activated the supraorbital nerve to evoke a blink reflex (BR), recorded via electromyography of the orbicularis oculi muscles. An active recording electrode was positioned on the lower eyelid, halfway between the inner and outer edges of the orbit, while the reference electrode was placed on the ipsilateral temple. Surface electrodes were used to deliver supraorbital nerve stimuli, with the cathode placed over the supraorbital notch and the anode positioned 3 cm away along the course of the nerve on the ipsilateral forehead. Percutaneous stimuli lasting 0.2 ms were applied at irregular intervals of 25–30 seconds. The stimulus intensity was set at three times the R2 motor threshold. Amplification of the responses was performed using a Digitimer D360R-4 device (Digitimer, Welwyn Garden City, UK). The signal was filtered using a frequency range of 50 Hz to 3 kHz. Data acquisition was carried out using Signal software, version 7.02 (Cambridge Electronic Devices, Cambridge, UK), and recorded on a laptop computer. For analysis, EMG traces were rectified, and all traces contaminated by EMG artifacts were discarded (Figure 1). Latency and area-under-the-curve (area) of R1, ipsilateral R2 (R2), and contralateral R2 (R2c) responses have been analysed.

### 2.3. Procedures

#### 2.3.1. Experiment 1: assessing the effect of a vestibular prepulse on the blink reflex area

Based on available data on vestibular perceptual and nystagmic thresholds (Seemungal et al., 2004), we choose four accelerations values above the perceptual threshold (5, 10, 20, or 30 degrees/second^2^ respectively), ensuring an activation of the vestibular system in all subjects. Experiment 1 consisted of three different conditions: condition 1, measured the unconditioned blink reflex; condition 2, measured the vestibular-conditioned blink reflex; condition 3, rotating the chair without evoking a blink reflex. Four trials were collected for condition 1 and 3, whereas in condition 2, eight trials for each angular acceleration were collected, four for right-to-left rotations and four for left-to-right rotations respectively. A total of 40 trials were collected.

#### 2.3.2. Experiment 2: assessing the effect of the vestibulo-ocular reflex suppression on PPI

To control for the possibility that any inhibition noted in Experiment 1 was linked secondarily to activation of the oculomotor system, we performed the vestibular pre-pulse task whilst subjects suppressed the vestibulo-ocular reflex by fixating on a visual cue. Experiment 2 consisted of two sub-experiments, each with two conditions: in 2A, we recorded an unconditioned blink reflex and a vestibular-conditioned blink reflex; in 2B, we record an unconditioned blink reflex and a vestibular conditioned blink reflex, but for the entire duration of the experiment subjects were instructed to keep their eyes open while staring at a chair-mounted head referenced red laser light in the dark (Jacobson et al., 2012), which was projected on a curtain surrounding the rotating chair. In both Experiment 2A and 2B, there were four trials, and the chair was rotated in the dark (except for point light source where indicated) with an angular acceleration of 10 degrees/second^2^.

#### 2.3.3. Experiment 3: assessing the vestibular PPI in patients with bilateral vestibular dysfunction

To corroborate our findings from previous experiments, we performed the vestibular PPI task in two patients clinically diagnosed with a bilateral peripheral vestibular failure. These patients’ peripheral vestibular failure was confirmed via video head impulse testing, bithermal caloric irrigation as well as by perceptual and nystagmic thresholds testing. Then, these patients underwent the same procedures outlined in Experiment 2a. In addition, to prove that other sensory modalities were still able to evoke PPI in these patients, we recorded the blink reflex response with and without a preceding somatosensory stimulation of the median nerve at the wrist. The intensity of the pre-pulse was set at two times the individual somatosensory threshold, and two interstimulus intervals (ie. 120 ms and 300 ms) were tested.

In all experiments, conditions were randomised, and an inter-trial interval of at least 25 seconds was used to avoid any possible habituation. Subjects were constantly monitored in dark through an infrared camera for safety reasons and to ensure that they remain alert for the duration of entire experiment. After each rotation, the lights were turned on to avoid any sleepiness and/or dump any residual post-rotational vestibular signal.

#### 2.3.4. Data analysis and statistics

For the main experiment (experiment 1), an a-priori power analysis was conducted using G*Power version 3.1.9.6 (Faul et al., 2007) to determine the minimum sample size required to test the study hypothesis. We based our power analysis on the amount of inhibition observed in the literature by using other sensory modalities. Results indicated the required sample size to achieve 80% power for detecting a medium effect, at a significance criterion of α = .05, was N = 15 for fixed effects one-way analysis of variance (ANOVA) test. Thus, the obtained sample size of N = 15 is adequate to test the study hypothesis. This number is also in line with previous sample sizes used in this research field. The experiment was not powered to account for differences in laterality given that we did not predict any differences between rightward or leftward rotations. Data comparing rightward vs. leftward rotations were reported. Homogeneity of variances was confirmed via Levene’s test for equality of variances.

In Experiment 1, we measured latency and area-under-the-curve (area) of the R1, R2, and R2c components of the blink reflex in each single rectified trace in control and test trials. We then average raw data per subject and per condition. We compared the effect of rotations at different accelerations on the latency and area of the R1, R2, and R2c responses of the blink reflex by using a one-way ANOVA. Multiple-comparison correction for three one-way ANOVAs was performed and p < 0.0167 was considered statistically significant. Effect size was calculated by using Cohen’s f (0.10 = small effect size; 0.25 = medium effect size; 0.40 or higher = large effect size) (Cohen, 2013). In case of a statistically significant main effect, Bonferroni corrected post-hoc tests were performed to assess the effect of each angular acceleration on the R1, R2, and R2c area.

For Experiment 2, we normalised the amount of inhibition for each subject, and we averaged the result per subject and per condition. PPI is expressed as 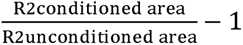. Similarly, modulation of R1 is expressed as 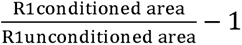. By using these formulas, values < 0 reflect an inhibition, whereas values > 0 reflects a facilitation of the response. We also reported the percentage of facilitation/inhibition by multiplying this value for 100. We then compared the effects of the VOR suppression on PPI by using a paired sample t-test. Because of multiple t-tests, we applied Bonferroni’s correction resulting in a significant P-value of 0.05/3 = 0.0167. Effect size was calculated by using Cohen’s d (0.20 = small effect size; 0.5 = medium effect size; 0.8 or higher = large effect size) (Cohen, 2013).

All statistical analyses were performed in SPSS version 26.

## 3. Results

All participants completed the experiment without reporting any adverse effect.

### 3.1. Experiment 1

Data for Experiment 1 are reported in Table 1 and 2, and in Figure 2 and 4. A one-way ANOVA failed to reveal any differences in the latencies of R1, R2, and R2c responses of the blink reflex between conditions (for R1: F(4, 70) = 0.071, p = 0.991, Cohen’s f = 0.004; for R2: F(4,70) = 0.091, p = 0.985, Cohen’s f = 0.005 ; for R2c: F(4,70) = 0.027, p = 0.999, Cohen’s f = 0.002) (see Table 1).

**Figure 2:**
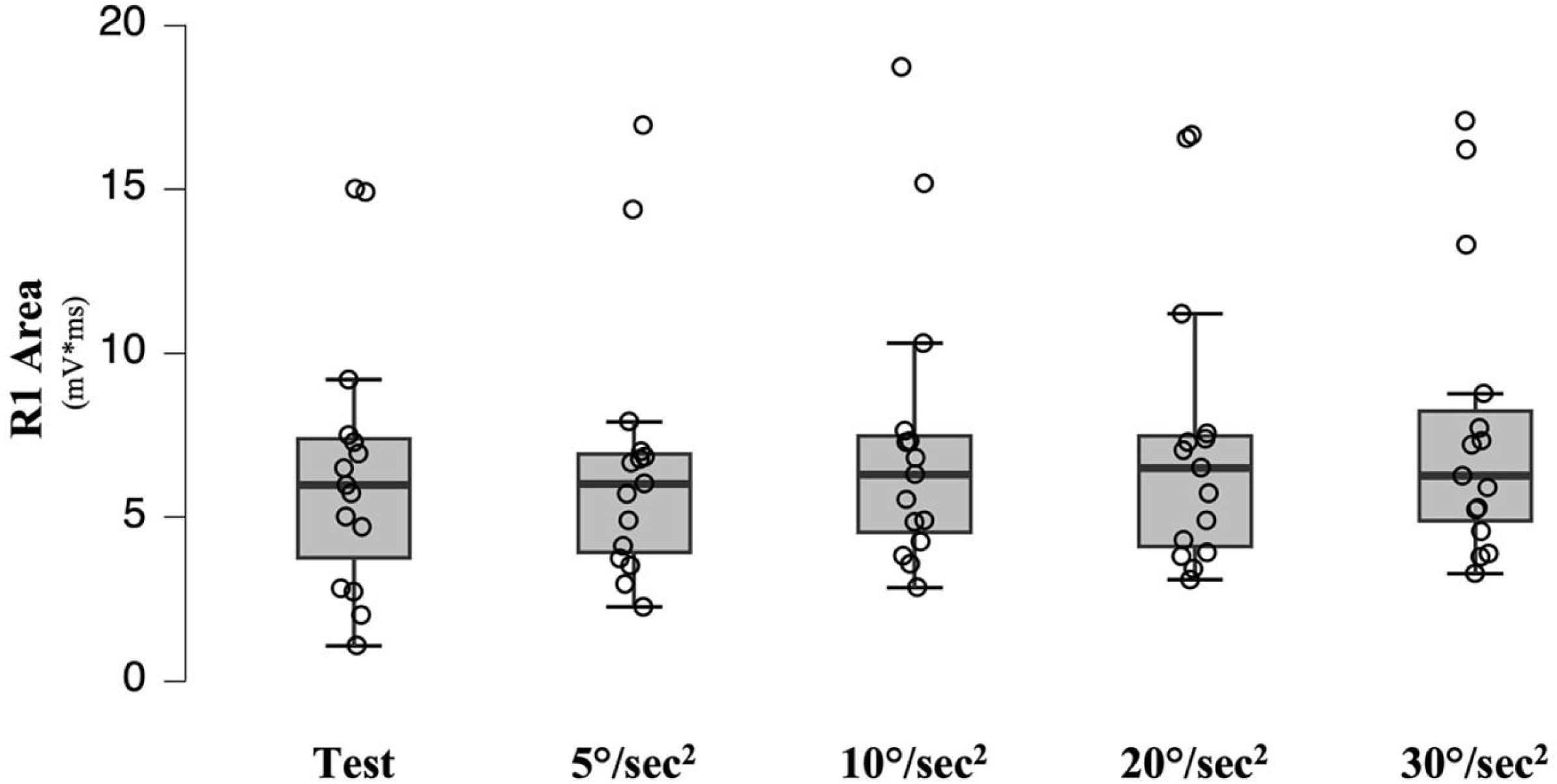
modulation of R1 area according to the magnitude of vestibular prepulse. R1 area (mV*ms) is presented for each condition. No statistically significant difference was observed, although a trend towards an increase in the area was noted.

**Table 1:** effect of a vestibular prepulse on the latency of the blink reflexes responses. Average, confidence interval and standard error for latencies (ms) of R1, R2, and R2c response are reported for each condition.

|  |  | 95% Confidence Interval Mean |  |  | Std. Error |
| --- | --- | --- | --- | --- | --- |
|  |  | Mean | Upper | Lower |  |
| **R1** | Test | 9.873 | 10.509 | 9.237 | 0.296 |
|  | 5°/sec<sup>2</sup> | 10.036 | 10.742 | 9.331 | 0.329 |
|  | 10°/sec<sup>2</sup> | 10.103 | 10.841 | 9.364 | 0.344 |
|  | 20°/sec<sup>2</sup> | 9.953 | 10.651 | 9.255 | 0.325 |
|  | 30°/sec<sup>2</sup> | 10.000 | 10.709 | 9.290 | 0.3307 |
| **R2** | Test | 28.413 | 30.101 | 26.72 | 0.787 |
|  | 5°/sec<sup>2</sup> | 28.763 | 29.970 | 27.555 | 0.562 |
|  | 10°/sec<sup>2</sup> | 28.764 | 30.197 | 27.329 | 0.668 |
|  | 20°/sec<sup>2</sup> | 28.890 | 30.284 | 27.495 | 0.650 |
|  | 30°/sec<sup>2</sup> | 28.926 | 30.366 | 27.487 | 0.671 |
| **R2c** | Test | 31.673 | 33.410 | 29.935 | 0.810 |
|  | 5°/sec<sup>2</sup> | 31.440 | 32.693 | 30.186 | 0.584 |
|  | 10°/sec<sup>2</sup> | 31.490 | 32.976 | 30.003 | 0.693 |
|  | 20°/sec<sup>2</sup> | 31.633 | 32.777 | 30.062 | 0.632 |
|  | 30°/sec<sup>2</sup> | 31.531 | 33.211 | 30.055 | 0.735 |

Figure 2 shows that there was no difference in the R1 area between conditions (F(4, 130) = 6.592, p = 0.837; Cohen’s f = 0.01), although a trend towards a facilitation was observed (Table 2).

**Table 2:** effect of a vestibular prepulse on the area of blink reflexes responses. Average, confidence interval and standard error for area (mV*ms) of R1, R2, and R2c responses are reported for each condition. Blink Reflex (BR) modulation is reported as a percentage, with facilitation represented by positive values and inhibition by negative values. P-values after Bonferroni correction are reported.

|  |  | 95% Confidence Interval Mean |  |  | Std. Error | BR modulation | p-value |
| --- | --- | --- | --- | --- | --- | --- | --- |
|  |  | Mean | Upper | Lower |  |  |  |
| **R1** | Test | 6.493 | 8.570 | 4.415 | 1.061 | - | - |
|  | 5°/sec<sup>2</sup> | 6.651 | 8.701 | 4.601 | 0.746 | +2.1% | n.s. |
|  | 10°/sec<sup>2</sup> | 7.287 | 9.522 | 5.051 | 0.797 | +11.9% | n.s. |
|  | 20°/sec<sup>2</sup> | 7.288 | 9.481 | 5.095 | 0.794 | +11.9% | n.s. |
|  | 30°/sec<sup>2</sup> | 7.719 | 9.940 | 5.497 | 0.806 | +18.5% | n.s. |
| **R2** | Test | 26.136 | 27.691 | 24.581 | 0.794 | - | - |
|  | 5°/sec<sup>2</sup> | 19.408 | 21.447 | 17.369 | 0.856 | -25.7% | P < 0.001 |
|  | 10°/sec<sup>2</sup> | 17.681 | 18.975 | 16.386 | 0.594 | -32.3% | P < 0.001 |
|  | 20°/sec<sup>2</sup> | 17.887 | 19.883 | 15.892 | 0.733 | -31.5% | P < 0.001 |
|  | 30°/sec<sup>2</sup> | 19.584 | 21.793 | 17.375 | 0.861 | -25.1% | P < 0.001 |
| **R2c** | Test | 21.944 | 24.857 | 19.030 | 1.487 | - | - |
|  | 5°/sec<sup>2</sup> | 16.464 | 18.513 | 14.414 | 0.954 | -24.9% | P < 0.05 |
|  | 10°/sec<sup>2</sup> | 15.216 | 17.283 | 13.148 | 0.814 | -31.7% | P < 0.001 |
|  | 20°/sec<sup>2</sup> | 15.209 | 16.889 | 13.528 | 0.654 | -31.7% | P < 0.001 |
|  | 30°/sec<sup>2</sup> | 17.711 | 20.114 | 15.308 | 1.089 | -19.3% | n.s. |

Figure 3 shows that the R2 area was different between conditions (F(4, 130) = 12.470, p < 0.0001; Cohen’s f = 0.29). The mean R2 area was different between the test and all the post-rotational responses (Table 2)

**Figure 3:**
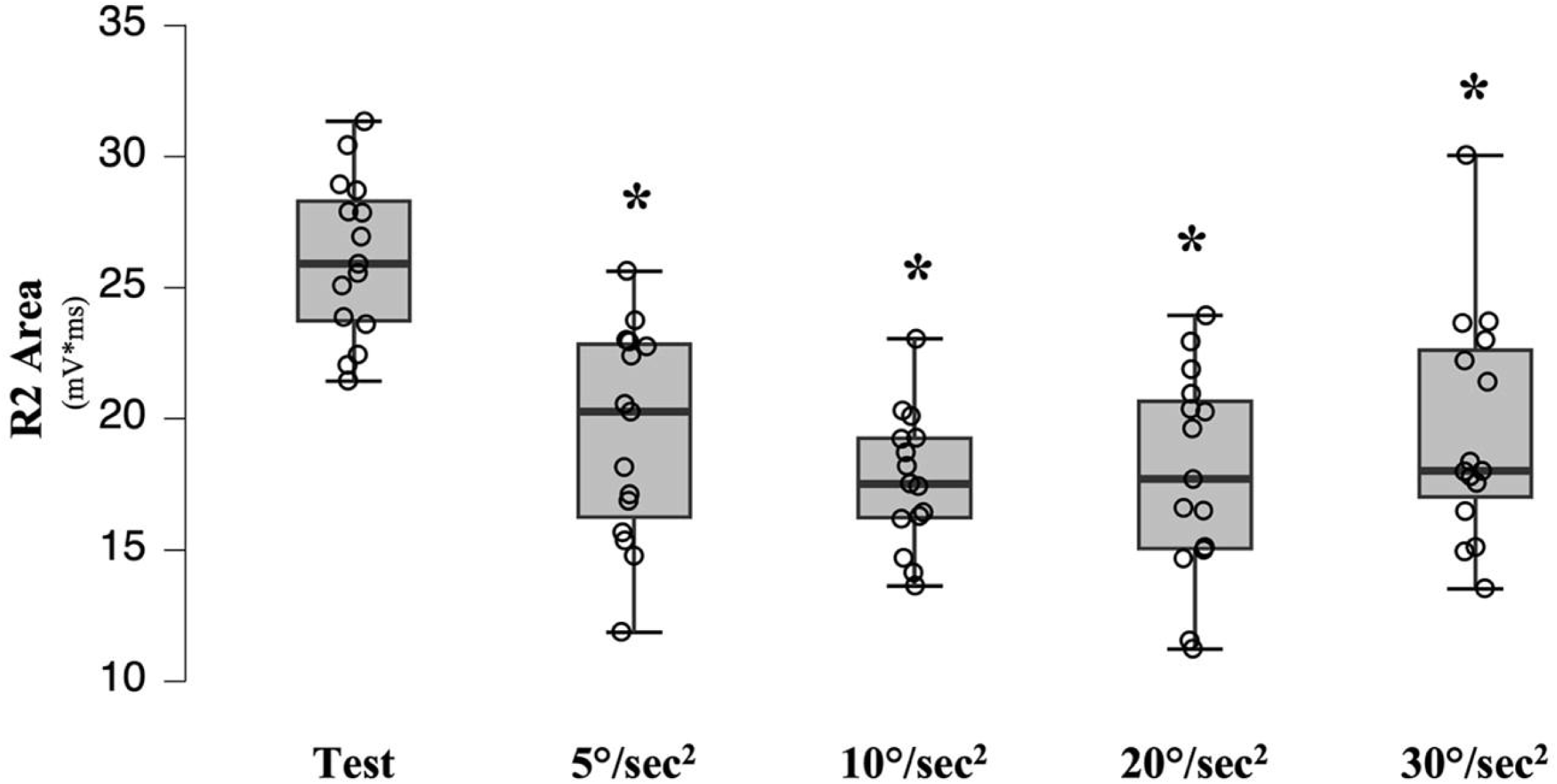
modulation of R2 area according to the magnitude of vestibular prepulse. R2 area (mV*ms) is presented for each condition. All vestibular prepulses reduced the R2 area of a subsequent test response. * < 0.05.

Similar results (F(4, 130) = 5.885, p < 0.0001; Cohen’s f = 0.15) were observed in the R2c response at 5°/s^2^, 10°/s^2^, and 20°/s^2^, while no difference was observed at 30°/s^2^ (Table 2). No difference between right and left rotations was observed for R1 (F(2, 130) = 0.219, p > 0.05, Cohen’s f = 0.063), R2 (F(2,130) = 22.330, p > 0.05, Cohen’s f = 0.261), and R2c (F(2,130) = 12.201, p > 0.05, Cohen’s f = 0.157).

### 3.2. Experiment 2

Data from Experiment 2A and 2B are reported in Table 3 and Figure 4.

**Table 3:** effect of the vestibulo-ocular reflex suppression on vestibular PPI. Average, confidence interval and standard error for area (mV*ms) of R1, R2, and R2c response are reported for each condition. PPI is reported as a percentage, with facilitation represented by positive values and inhibition by negative values.

|  |  | 95% Confidence<br>Interval Mean |  |  |  |  |  |  |
| --- | --- | --- | --- | --- | --- | --- | --- | --- |
|  | VOR |  | Mean | Upper | Lower | Std. Error | PPI | p-value |
| R1 | Present | Test | 3.658 | 4.262 | 3.054 | 0.308 | - | - |
| R1 |  | 10°/sec2 | 4.498 | 5.254 | 3.743 | 0.385 | +23% | - |
| R1 | Suppressed | Test | 3.537 | 4.246 | 2.828 | 0.362 | - | - |
| R1 |  | 10°/sec2 | 4.351 | 5.115 | 3.587 | 0.390 | +23% | 0.438 |
| R2 | Present | Test | 22.891 | 24.896 | 20.886 | 1.023 | - | - |
| R2 |  | 10°/sec2 | 17.359 | 19.024 | 15.694 | 0.849 | -24.2% | - |
| R2 | Suppressed | Test | 20.798 | 22.588 | 19.007 | 0.914 | - | - |
| R2 |  | 10°/sec2 | 15.607 | 17.461 | 13.752 | 0.946 | -25% | 0.732 |
| R2c | Present | Test | 20.888 | 22.953 | 18.824 | 1.053 | - | - |
| R2c |  | 10°/sec2 | 16.239 | 17.748 | 14.731 | 0.769 | -22.3% | - |
| R2c | Suppressed | Test | 18.753 | 20.463 | 17.042 | 0.873 | - | - |
| R2c |  | 10°/sec2 | 14.836 | 16.727 | 12.944 | 0.965 | -21.9% | 0.930 |

**Figure 4:**
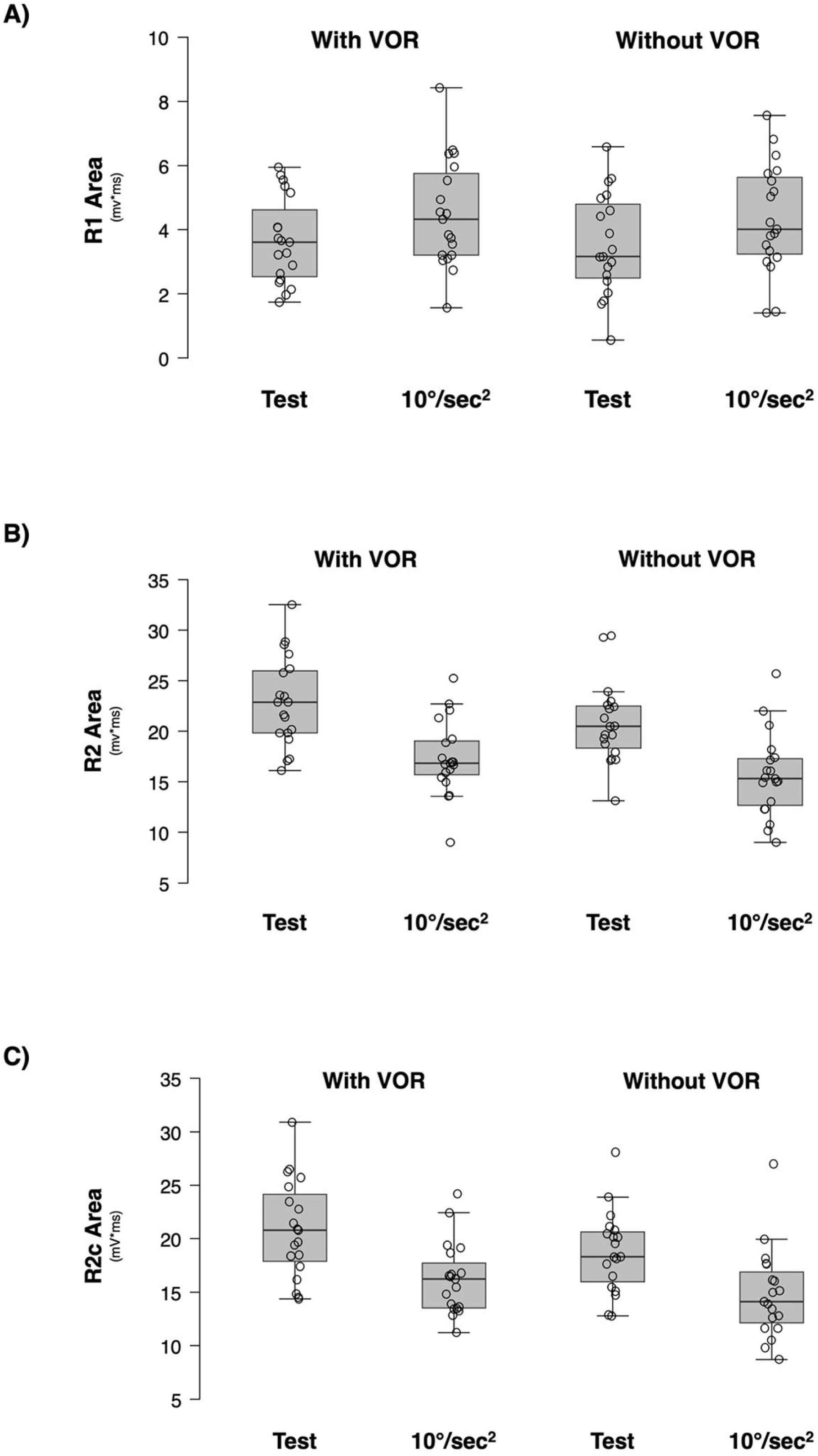
modulation of R1 (A), R2 (B), and R2c (C) area with and without suppression of the vestibulo-ocular reflex. Grand average and standard error for trials with and without vestibulo-ocular reflex are presented for R1, R2, and R2c area (mV*ms). Vestibulo-ocular reflex suppression did not change the magnitude of PPI. VOR: vestibulo-ocular reflex.

We compared the amount of inhibition obtained with and without vestibulo-ocular reflex suppression. A paired sample t-test did not show any difference in the R1 (t = -0.793; p > 0.05; Cohen’s d = 0.226), R2 (t = 0.348; p > 0.05; Cohen’s d = 0.075), and R2c (t = -0.089; p > 0.05; Cohen’s d = 0.034) responses between conditions.

To explore a possible gender-related modulation of PPI, we compared the magnitude of PPI between males (n=12) and females (n=10). No statistically significant difference has been noted in the R1, R2, and R2c area (see table 4), although a slightly higher inhibition is observed in males.

**Table 4:**
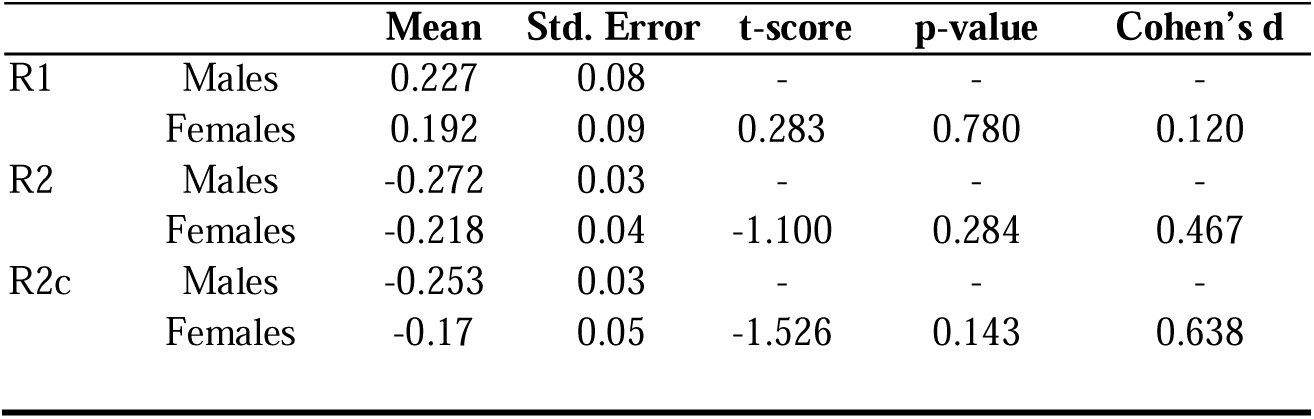
effect of gender on vestibular PPI. Magnitude of inhibition and standard error of R1, R2, and R2c responses are reported for each group. Facilitation is represented by positive values and inhibition by negative values. T scores, p-values, and Cohen’s d are reported.

### 3.3. Experiment 3

Results from experiment 3 are reported in Figure 5 and Table 5. Briefly, both patients did not show any inhibition when the vestibular pre-pulse was delivered. PPI was conserved for the somatosensory modality at both interstimulus intervals.

**Figure 5:**
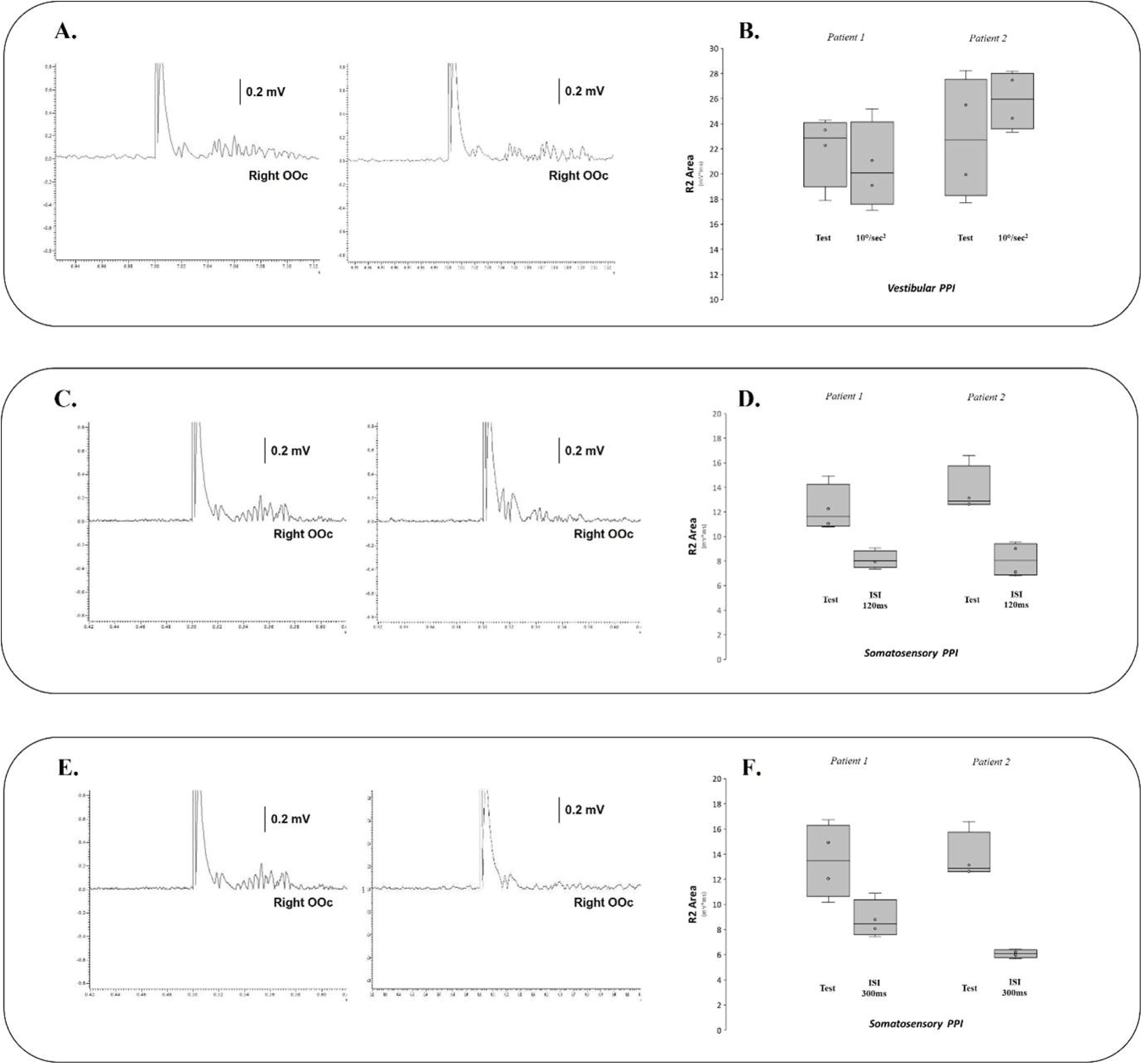
modulation of the R2 component of the blink reflex in patients with bilateral vestibular failure. In the first panel, in A the unconditioned R2 response (left) and the R2 response conditioned by a vestibular prepulse (right). In B, average and standard deviation of the R2 response area for trials unconditioned and conditioned by a vestibular prepulse. No inhibition is present. In the second panel, in C the unconditioned R2 response (left) and the R2 response conditioned by a somatosensory prepulse delivered 120 ms before the blink reflex (right). In D, the corresponding average and standard deviation of the R2 response area for trials unconditioned and conditioned by a somatosensory prepulse. Somatosensory PPI is preserved. Similarly, In the third panel, in E the unconditioned R2 response (left) and the R2 response conditioned by a somatosensory prepulse delivered 300 ms before the blink reflex (right). In F, the corresponding average and standard deviation of the R2 response area for trials unconditioned and conditioned by a somatosensory prepulse. Somatosensory PPI is again preserved. Area: mV*ms. OOc: orbiculary oculi muscle. ISI: interstimuls interval. mV: millivolts.

**Table 5:** prepulse inhibition in patients with bilateral vestibular failure. Magnitude of inhibition/facilitation of R1, R2, and R2c responses are reported for each patient for each prepulse modality. Facilitation is represented by positive values and inhibition by negative values. vPPI: vestibular prepulse inhibition; sPPI 120 ms: somatosensory prepulse inhibition with an interstimulus interval of 120 ms. sPPI 300 ms: somatosensory prepulse inhibition with an interstimulus interval of 300 ms.

|  |  | **vPPI** | **sPPI 120 ms** | **sPPI 300 ms** |
| --- | --- | --- | --- | --- |
|  |  | **%** | **%** | **%** |
| R1 | Patient 1 | -9.44 | +14.65 | +8.5 |
|  | Patient 2 | -4.95 | +108 | +15.25 |
| R2 | Patient 1 | -6.25 | -33.73 | -34.69 |
|  | Patient 2 | +13.16 | -40.97 | -55.75 |
| R2c | Patient 1 | -8.11 | -57.12 | -29.66 |
|  | Patient 2 | +20.4 | -40.89 | -60.85 |

## 4. Discussion

The present data demonstrate how the activation of the vestibular system can inhibit the R2 response of a subsequent blink reflex. Our data are the first that support the notion for a vestibular-evoked gating mechanism, and this can be quantified using a standard measure – the PPI-as previously demonstrated for other prepulse modalities. Our data did not show any modulation for increasing vestibular angular accelerations, however the range we used was clustered toward the lower end of head accelerations found in the ecological movements. Notably, the lack of effect by vestibulo-ocular reflex suppression and the lack of inhibition in patients with bilateral vestibular failure indicate that our data were linked to vestibular activation specifically (and not an indirect effect of eye movements and the associated motor circuits or a direct effect of a proprioceptive modulation).

### 4.1. Vestibular PPI and Functional neuroanatomy

The finding that the activation of the vestibular system modulates the blink reflex supports the hypothesis that a vestibular pulse can activate the PPI network. PPI is primarily regulated in the brainstem, and previous studies have proposed an anatomical network for somatosensory and auditory prepulses, which involves fibres in the superior colliculus and in the lower auditory system, respectively (Gomez-Nieto et al., 2020, Rohleder et al., 2016). Both networks converge in the PPN area, which then regulates the final motor outcome through the nucleus reticularis pontis caudalis (Gomez-Nieto et al., 2020). Visual and auditory inputs were control for by testing in a silent and dark room with continuous background noise of 30 dB arising primarly from electrical hum of the laboratory equipment. Somatosensory inputs were minimised using a vibrationless computer-controlled rotating chair. In concert with the results from the vestibular failure patients, our data suggest that the inhibition of the blink reflex solely depends on pre-activation of the vestibular system, as seen in classic prepulse inhibition paradigms (Valls-Sole et al., 1999). The vestibular activation was carefully controlled by the use of 2 seconds long, symmetric triangular stimuli, so as to avoid any post-rotational vestibular activation linked to prolonged displacement of the cupola (Guedry, 1974). This ensures that the vestibular activation does not extend beyond the duration of the chair rotation, thereby minimising any residual activity in the peripheral vestibular system while evoking the blink reflex.

The vestibular network shares several structures with the PPI functional network, including the superior colliculus and the PPN (Kirsch et al., 2016). Therefore, it is likely that a vestibular input is integrated in the PPI network, inducing an inhibition that can be measured by measuring the area of the R2 response of the blink reflex. Interestingly, vPPI is independent from the presence of the vestibulo-ocular reflex. Previous studies have shown a differential integration and modulation of vestibular inputs subserving different functions, e.g. those circuits processing the vestibulo-ocular reflex versus vestibular perceptual signals of self-motion (Seemungal, 2015). Interestingly PPN DBS modulates both vestibular perception and postural control (Yousif et al., 2016). It follows that assessing links between the vPPI and a variety of vestibular-mediated functions (perception and/or balance) would be of potential clinical utility, particularly since recent studies show overlap between higher order brain circuits – themselves modulated by the PPN (Inagaki et al., 2022) - mediating balance and vestibular perception (Calzolari et al., 2021). In addition, our results are consistent with the observation that if the intensity of the prepulse stimulus is enough to generate a response (i.e. the VOR in our experiment), such response does not interfere with inhibition of the R2 response (Kofler et al., 2023). Therefore, it is not surprising that vPPI is independent form the activation of the VOR network. Further experiments will be needed to elucidate the relationship between vPPI and vestibular motion perception, particularly with respect to individual vestibular perceptual thresholds.

From a neuroanatomical perspective, the PPN is one of the midbrain structures where the circuits that mediate PPI overlap with the vestibular network (Kirsch et al., 2016). The PPN is the major component of the mesencephalic locomotor area, located on the caudal mesencephalic region, and it participates in controlling several functions, although it has been mainly related to locomotion, especially in animal models. However, in humans, it seems to be involved in a broader set of functions, such as posture (Breit et al., 2023, Joza et al., 2022, Lin et al., 2023), eye movements (Ewenczyk et al., 2017, Gallea et al., 2021), sleep (Gallea et al., 2017), and cognition (Mena-Segovia and Bolam, 2011, Petzold et al., 2015). Some of these functions rely on the integration of different sensory feedback loops, and PPI has been already advocated as a promising tool to measure how the brainstem integrates peripheral feedbacks (Kofler et al., 2023), also during tasks where the modulation of the PPN may have a pivotal role (Versace et al., 2019). Indeed, we showed here how vestibular inputs can be integrated at the level of the brainstem similarly to other sensory modalities (i.e. somatosensory and auditory). As vestibular inputs are relevant for most of the activities mediated by the PPN, such as gait, posture, and eye movements (Cullen, 2023), we suggest that vestibular PPI might be a useful tool to measure how the vestibular signal is integrated with respects to the abovementioned PPN-related activities.

Our data cannot elucidate whether a direct connection between the vestibular nuclei and the PPN is responsible for the observed PPI. Indirect connections between the vestibular nuclei, the cerebellum, and the PPN can be responsible for the activation of the PPN (Kirsch et al., 2016), as well as other cortical structures may be responsible for this modulation. In fact, while the primary neural circuitry responsible for mediating PPI is predominantly located in the brainstem, and PPI is typically considered an automatic process (Lei et al., 2021), it is important to note that PPI can also be influenced by higher-order cognitive processes, including attention and emotion (Lei et al., 2021), and that thalamic, striatal and frontal lobe activation has also been observed during PPI in healthy groups (Fendt et al., 2001, Naysmith et al., 2021, Rohleder et al., 2016, Santos-Carrasco and De la Casa, 2023, Schmajuk and Larrauri, 2005). Further research is needed to integrate the vestibular network with the PPI functional network and to rule out whether a direct connection between the vestibular nuclei and the PPN exists in humans.

### 4.2. Sensitivity of vestibular PPI

Similar to other sensory modalities, we were expecting for vestibular PPI to be reactive to the intensity of the prepulse and to the gender of the participant. Interestingly however, the inhibition was present at all the values of angular acceleration tested, without a clear angular acceleration-related modulation. This contrasts with previous findings in auditory and somatosensory PPI studies (Blumenthal, 2015), where increasing the amplitude of the prepulse resulted in greater inhibition. However, although we selected four pulses of angular acceleration which were 5 to 30 times larger than the perceptual threshold reported in healthy subjects, and 10 to 60 times larger than the vestibulo-ocular reflex (nystagmic) threshold (Seemungal et al., 2004), these values are many times less than typical head peak head angular accelerations during normal movements (Carriot et al., 2014). It is likely that the lack of modulation relates to the use of stimuli at the bottom end of the ecological dynamic range of the vestibular apparatus, although additional studies will be required to confirm the intensity-dependent vPPI modulation with much wider range of angular head accelerations (Carriot et al., 2014).

With regards to the gender, we noticed a trend towards a difference in PPI, with a higher magnitude of inhibition in men compare to women. This is in line with previous literature, which reports a difference of approximately 20-40% in the magnitude of PPI between healthy young men and women (Aasen et al., 2005, Abel et al., 1998, Swerdlow et al., 1993). This difference might be even more consistent when considering the menstrual cycle status, as more inhibition has been described in the early follicular phase compared to the luteal phase (Kumari et al., 2010), and no differences in PPI have been noticed between men and women on hormonal contraception (Naysmith et al., 2022). In our study, we did not record data on participants’ menstrual cycle or use of oral contraception. Future studies are needed to address this possible biological difference.

### 4.3. Technical Considerations

Finally, there are two relevant considerations needed on the prepulse and the interstimulus interval we used for this experiment. Firstly, our prepulse lasts longer than the ones used in other PPI studies using different modalities. It is possible that by using a 2-second long prepulse we might have reduced the magnitude of the subsequent inhibition, as PPI is sensitive to variations in prepulse duration (Blumenthal, 2015). Secondly, we only investigated one interstimulus interval (i.e. 300 ms). We choose this interstimulus interval as it has been shown to give a reliable inhibition in other PPI studies (Valls-Sole et al., 1999). However, it is possible for shorter interstimulus interval to be more effective in inducing an inhibition in the blink reflex. This would also support the existence of a direct connection between the vestibular nuclei and the PPN, ensuring a more rapid filtering of the vestibular prepulse in the sensorimotor gating network. Therefore, future studies might be focused on investigating others interstimulus intervals, both to explore which is the interstimulus interval with the strongest inhibition and whether at longer interstimulus intervals facilitation is present.

### 4.4. Clinical Implications

The modulation of vestibular signalling by means of non-invasive brain stimulation techniques has gained increasing interest over the recent years as a treatment for some neurodegenerative conditions. Particularly, galvanic vestibular stimulation has been used to improve balance and vestibular outputs in patients with bilateral vestibulopathy (Schniepp et al., 2018), Parkinson’s disease (Mahmud et al., 2022), multiple sclerosis (Lotfi et al., 2021), and dizziness (Woll et al., 2019).

Galvanic vestibular stimulation may modulate balance, potentially by modulating the PPN-thalamic connections, as shown in neuroimaging studies (Cai et al., 2018). However, a proper neurophysiological measure of the interaction between the vestibular system and PPN function is missing. To this extent, vestibular PPI is a promising tool, as it offers the opportunity to investigate the integration of vestibular inputs in the PPN. Further studies will be needed to elucidate the neuroanatomy and neurophysiology of vestibular PPI, as these are vital to ascertaining whether this novel technique has any potential as a marker of vestibular-motor integration at the brainstem level and hence whether it can be used to interrogate the utility of treatments for improving PPN-mediated functions such as balance in human brain disease.

## 5. Conclusion

Our findings support the existence of a vestibular PPI in humans, which can be measured by means of the blink reflex. A vestibular prepulse should be added to the list, alongside the auditory and somatosensory prepulses, as capable of evoking an inhibition in the blink reflex response. Vestibular PPI might reflect the sensorimotor gating of vestibular inputs at the level of the brainstem as well as being a measure of how the PPN integrates vestibular information. Further studies will confirm the role of vestibular PPI as a possible marker of vestibular-motor integration at the brainstem level, particularly in relation to non-invasive brain stimulation techniques of the vestibular system.

## References

1. Aasen I, Kolli L, Kumari V. Sex effects in prepulse inhibition and facilitation of the acoustic startle response: implications for pharmacological and treatment studies. J Psychopharmacol 2005;19(1):39–45.

2. Abel K, Waikar M, Pedro B, Hemsley D, Geyer M. Repeated testing of prepulse inhibition and habituation of the startle reflex: a study in healthy human controls. J Psychopharmacol 1998;12(4):330–7.

3. Aravamuthan BR, Angelaki DE. Vestibular responses in the macaque pedunculopontine nucleus and central mesencephalic reticular formation. Neuroscience 2012;223:183–99.

4. Blumenthal TD. Presidential Address 2014: The more-or-less interrupting effects of the startle response. Psychophysiology 2015;52(11):1417–31.

5. Breit S, Milosevic L, Naros G, Cebi I, Weiss D, Gharabaghi A. Structural-Functional Correlates of Response to Pedunculopontine Stimulation in a Randomized Clinical Trial for Axial Symptoms of Parkinson’s Disease. J Parkinsons Dis 2023;13(4):563–73.

6. Cai J, Lee S, Ba F, Garg S, Kim LJ, Liu A, et al. Galvanic Vestibular Stimulation (GVS) Augments Deficient Pedunculopontine Nucleus (PPN) Connectivity in Mild Parkinson’s Disease: fMRI Effects of Different Stimuli. Front Neurosci 2018;12:101.

7. Calzolari E, Chepisheva M, Smith RM, Mahmud M, Hellyer PJ, Tahtis V, et al. Vestibular agnosia in traumatic brain injury and its link to imbalance. Brain 2021;144(1):128–43.

8. Carriot J, Jamali M, Chacron MJ, Cullen KE. Statistics of the vestibular input experienced during natural self-motion: implications for neural processing. J Neurosci 2014;34(24):8347–57.

9. Cohen J. Statistical power analysis for the behavioral sciences: Academic press, 2013.

10. Csomor PA, Vollenweider FX, Feldon J, Yee BK. On the feasibility to detect and to quantify prepulse-elicited reaction in prepulse inhibition of the acoustic startle reflex in humans. Behav Brain Res 2005;162(2):256–63.

11. Cullen KE. Vestibular motor control. Handb Clin Neurol 2023;195:31–54.

12. Ewenczyk C, Mesmoudi S, Gallea C, Welter ML, Gaymard B, Demain A, et al. Antisaccades in Parkinson disease: A new marker of postural control? Neurology 2017;88(9):853–61.

13. Faul F, Erdfelder E, Lang AG, Buchner A. G*Power 3: a flexible statistical power analysis program for the social, behavioral, and biomedical sciences. Behav Res Methods 2007;39(2):175–91.

14. Fendt M, Li L, Yeomans JS. Brain stem circuits mediating prepulse inhibition of the startle reflex. Psychopharmacology (Berl) 2001;156(2-3):216–24.

15. Gallea C, Ewenczyk C, Degos B, Welter ML, Grabli D, Leu-Semenescu S, et al. Pedunculopontine network dysfunction in Parkinson’s disease with postural control and sleep disorders. Mov Disord 2017;32(5):693–704.

16. Gallea C, Wicki B, Ewenczyk C, Rivaud-Pechoux S, Yahia-Cherif L, Pouget P, et al. Antisaccade, a predictive marker for freezing of gait in Parkinson’s disease and gait/gaze network connectivity. Brain 2021;144(2):504–14.

17. Garcia-Rill E, Saper CB, Rye DB, Kofler M, Nonnekes J, Lozano A, et al. Focus on the pedunculopontine nucleus. Consensus review from the May 2018 brainstem society meeting in Washington, DC, USA. Clin Neurophysiol 2019;130(6):925–40.

18. Gomez-Nieto R, Hormigo S, Lopez DE. Prepulse Inhibition of the Auditory Startle Reflex Assessment as a Hallmark of Brainstem Sensorimotor Gating Mechanisms. Brain Sci 2020;10(9).

19. Guedry F. Psychophysics of Vestibular Sensation. In: Kornhuber, HH (eds) Vestibular System Part 2: Psychophysics, Applied Aspects and General Interpretations Handbook of Sensory Physiology 1974;vol 6 / 2. Springer, Berlin, Heidelberg.

20. Hanzlikova Z, Kofler M, Slovak M, Vechetova G, Fecikova A, Kemlink D, et al. Prepulse inhibition of the blink reflex is abnormal in functional movement disorders. Mov Disord 2019;34(7):1022–30.

21. Horowitz SS, Blanchard J, Morin LP. Medial vestibular connections with the hypocretin (orexin) system. J Comp Neurol 2005;487(2):127–46.

22. Inagaki HK, Chen S, Ridder MC, Sah P, Li N, Yang Z, et al. A midbrain-thalamus-cortex circuit reorganizes cortical dynamics to initiate movement. Cell 2022;185(6):1065–81 e23.

23. Jacobson GP, Piker EG, Do C, McCaslin DL, Hood L. Suppression of the vestibulo-ocular reflex using visual and nonvisual stimuli. Am J Audiol 2012;21(2):226–31.

24. Joza S, Camicioli R, Martin WRW, Wieler M, Gee M, Ba F. Pedunculopontine Nucleus Dysconnectivity Correlates With Gait Impairment in Parkinson’s Disease: An Exploratory Study. Front Aging Neurosci 2022;14:874692.

25. Karachi C, Grabli D, Bernard FA, Tande D, Wattiez N, Belaid H, et al. Cholinergic mesencephalic neurons are involved in gait and postural disorders in Parkinson disease. J Clin Invest 2010;120(8):2745–54.

26. Kirsch V, Keeser D, Hergenroeder T, Erat O, Ertl-Wagner B, Brandt T, Dieterich M. Structural and functional connectivity mapping of the vestibular circuitry from human brainstem to cortex. Brain Struct Funct 2016;221(3):1291–308.

27. Kofler M, Valls-Sole J, Thurner M, Pucks-Faes E, Versace V. In the spotlight: How the brainstem modulates information flow. Clin Neurophysiol 2023;148:52–64.

28. Kumari V, Konstantinou J, Papadopoulos A, Aasen I, Poon L, Halari R, Cleare AJ. Evidence for a role of progesterone in menstrual cycle-related variability in prepulse inhibition in healthy young women. Neuropsychopharmacology 2010;35(4):929–37.

29. Lei M, Ding Y, Meng Q. Neural Correlates of Attentional Modulation of Prepulse Inhibition. Front Hum Neurosci 2021;15:649566.

30. Lin C, Ridder MC, Sah P. The PPN and motor control: Preclinical studies to deep brain stimulation for Parkinson’s disease. Front Neural Circuits 2023;17:1095441.

31. Lotfi Y, Farahani A, Azimiyan M, Moossavi A, Bakhshi E. Comparison of efficacy of vestibular rehabilitation and noisy galvanic vestibular stimulation to improve dizziness and balance in patients with multiple sclerosis. J Vestib Res 2021;31(6):541–51.

32. Mahmud M, Hadi Z, Prendergast M, Ciocca M, Saad AR, Pondeca Y, et al. The effect of galvanic vestibular stimulation on postural balance in Parkinson’s disease: A systematic review and meta-analysis. J Neurol Sci 2022;442:120414.

33. Mena-Segovia J, Bolam JP. Phasic modulation of cortical high-frequency oscillations by pedunculopontine neurons. Prog Brain Res 2011;193:85–92.

34. Millian-Morell L, Lopez-Alburquerque T, Rodriguez-Rodriguez A, Gomez-Nieto R, Carro J, Meilan JJG, et al. Relations between Sensorimotor Integration and Speech Disorders in Parkinson’s Disease. Curr Alzheimer Res 2018;15(2):149–56.

35. Naysmith LF, Kumari V, Williams SCR. Neural mapping of prepulse-induced startle reflex modulation as indices of sensory information processing in healthy and clinical populations: A systematic review. Hum Brain Mapp 2021;42(16):5495–518.

36. Naysmith LF, Williams SCR, Kumari V. The influence of stimulus onset asynchrony, task order, sex and hormonal contraception on prepulse inhibition and prepulse facilitation: Methodological considerations for drug and imaging research. J Psychopharmacol 2022;36(11):1234–42.

37. Petzold A, Valencia M, Pal B, Mena-Segovia J. Decoding brain state transitions in the pedunculopontine nucleus: cooperative phasic and tonic mechanisms. Front Neural Circuits 2015;9:68.

38. Rohleder C, Wiedermann D, Neumaier B, Drzezga A, Timmermann L, Graf R, et al. The Functional Networks of Prepulse Inhibition: Neuronal Connectivity Analysis Based on FDG-PET in Awake and Unrestrained Rats. Front Behav Neurosci 2016;10:148.

39. Santos-Carrasco D, De la Casa LG. Prepulse inhibition deficit as a transdiagnostic process in neuropsychiatric disorders: a systematic review. BMC Psychol 2023;11(1):226.

40. Schmajuk NA, Larrauri JA. Neural network model of prepulse inhibition. Behav Neurosci 2005;119(6):1546–62.

41. Schniepp R, Boerner JC, Decker J, Jahn K, Brandt T, Wuehr M. Noisy vestibular stimulation improves vestibulospinal function in patients with bilateral vestibulopathy. J Neurol 2018;265(Suppl 1):57–62.

42. Seemungal BM. The Components of Vestibular Cognition--Motion Versus Spatial Perception. Multisens Res 2015;28(5-6):507–24.

43. Seemungal BM, Gunaratne IA, Fleming IO, Gresty MA, Bronstein AM. Perceptual and nystagmic thresholds of vestibular function in yaw. J Vestib Res 2004;14(6):461–6.

44. Swerdlow NR, Auerbach P, Monroe SM, Hartston H, Geyer MA, Braff DL. Men are more inhibited than women by weak prepulses. Biol Psychiatry 1993;34(4):253–60.

45. Valls-Sole J, Munoz JE, Valldeoriola F. Abnormalities of prepulse inhibition do not depend on blink reflex excitability: a study in Parkinson’s disease and Huntington’s disease. Clin Neurophysiol 2004;115(7):1527–36.

46. Valls-Sole J, Valldeoriola F, Molinuevo JL, Cossu G, Nobbe F. Prepulse modulation of the startle reaction and the blink reflex in normal human subjects. Exp Brain Res 1999;129(1):49–56.

47. Versace V, Campostrini S, Sebastianelli L, Saltuari L, Valls-Sole J, Kofler M. Influence of posture on blink reflex prepulse inhibition induced by somatosensory inputs from upper and lower limbs. Gait Posture 2019;73:120–5.

48. Woll J, Sprenger A, Helmchen C. Postural control during galvanic vestibular stimulation in patients with persistent perceptual-postural dizziness. J Neurol 2019;266(5):1236–49.

49. Woolf NJ, Butcher LL. Cholinergic systems in the rat brain: IV. Descending projections of the pontomesencephalic tegmentum. Brain Res Bull 1989;23(6):519–40.

50. Yousif N, Bhatt H, Bain PG, Nandi D, Seemungal BM. The effect of pedunculopontine nucleus deep brain stimulation on postural sway and vestibular perception. Eur J Neurol 2016;23(3):668–70.

51. Zoetmulder M, Biernat HB, Nikolic M, Korbo L, Jennum PJ. Sensorimotor gating deficits in multiple system atrophy: comparison with Parkinson’s disease and idiopathic REM sleep behavior disorder. Parkinsonism Relat Disord 2014;20(3):297–302.

